# SEPepQuant enables comprehensive protein isoform characterization in shotgun proteomics

**DOI:** 10.1101/2022.11.03.515027

**Authors:** Yongchao Dou, Xinpei Yi, Lindsey K. Olsen, Bing Zhang

**Author notes:** Correspondence should be addressed to B.Z.

## Abstract

Shotgun proteomics is essential for protein identification and quantification in biomedical research, but protein isoform characterization is inhibited by the extensive number of peptides shared across proteins, hindering our understanding of protein isoform regulation and their roles in normal and disease biology. We systematically assess the challenge and opportunities of shotgun proteomics-based protein isoform characterization using *in silico* and experimental data, and then present SEPepQuant, a graph theory-based approach to enable isoform characterization. Using published data from one induced pluripotent stem cell study and two human hepatocellular carcinoma studies, we demonstrate the ability of SEPepQuant in addressing the key limitations of existing methods, providing more comprehensive proteome characterization, identifying hundreds of isoform-level regulation events, and facilitating streamlined cross-study comparisons. Our analysis provides solid evidence to support a widespread role of protein isoform regulation in normal and disease processes, and SEPepQuant has broad applications to biological and translational research.

## Introduction

Alternative splicing of precursor messenger RNA (pre-mRNA) is an essential post-transcriptional process that is believed to underly increased cellular and functional complexity in eukaryotic organisms^1,2^. This process is highly regulated, and dysregulated RNA splicing has been linked to a wide range of diseases such as retinal and developmental disorders, neurodegenerative diseases, and cancer^3,4^. High-throughput sequencing-based transcriptomic studies have showed that most human protein-coding genes undergo alternative splicing to produce multiple mRNA isoforms^5^. Mass spectrometry (MS)-based shotgun proteomics is the primary method for protein identification and quantification from biological samples^6^, but shotgun proteomics studies have provided very limited information on protein isoforms due to intrinsic challenges in data analysis. In fact, the extent to which transcript isoform complexity propagates to the proteome remains controversial^7-9^, and systematic investigation of the roles of protein isoforms in normal and disease biology is largely lacking^10^.

In a shotgun proteomics experiment, proteins extracted from biological samples are digested into peptides using enzymes such as trypsin and then analyzed by liquid chromatography-tandem mass spectrometry (LC-MS/MS). Each LC-MS/MS run generates thousands of spectra, which serve as the basis for the identification and quantification of peptides and proteins. Many bioinformatics tools have been developed to perform these essential computational tasks in shotgun proteomics data analysis, such as MaxQuant^11^, Trans-proteomic pipeline^12^, OpenMS^13^, FragPipe^14,15^, among others. Despite algorithmic and implementation differences, these tools share a similar workflow. First, observed MS/MS spectra are searched against a reference protein database for peptide identification. Next, identified peptides are used to infer a list of proteins that are assumed to be present in the sample, a process known as protein inference. Finally, inferred proteins are quantified based on the signal intensity measurements of the constituting peptides.

The difficulty in protein isoform characterization is tightly tied to the challenge in protein inference caused by the large number of degenerate peptides, which are peptides that can be mapped to multiple proteins due to a high level of sequence similarity between protein isoforms encoded by the same gene, or genes in the same gene family. The current best practice in the field is to collapse proteins with the same set of supporting peptides together with those that are supported by a subset of these peptides into a protein group^16,17^. For protein quantification, peptides shared by multiple protein groups are either ignored or assigned to the group with the largest number of associated peptides, and typically, one representative protein (i.e., the one with the largest number of associated peptides) is selected from each protein group for reporting and downstream analysis^11,14^. This parsimonious approach plays a critical role in preventing overstating the number of proteins in protein inference, however, it also limits the potential for protein isoform characterization. First, protein isoforms without uniquely identified peptides are essentially ignored. Secondly, the assignment of shared peptides to the protein groups and proteins with the largest number of associated peptides may not necessarily be the correct solution. Due to the hard-to-solve challenge in protein isoform discrimination, an alternative option is to perform gene group-based quantification, which is implemented in tools such as gpGrouper^18^ and FragPipe and used in some studies^19,20^. Because most gene groups only include a single gene, and the number of peptides shared between different gene groups is much smaller, gene group-based quantification greatly reduces uncertainty and potential misinterpretation of shared peptides. However, protein isoform information is lost completely with this approach.

In this study, we systematically assess the challenge and opportunities of shotgun proteomics-based protein isoform characterization using *in silico* digestion data and experimental data from a published induced pluripotent stem cell (iPSC) study^21^ and two published human hepatocellular carcinoma (HCC) studies^22,23^. We introduce a graph theory-based approach to accurately represent the peptide, protein, and gene relationships and to maximize protein isoform information that can be extracted from shotgun proteomics studies. Our method is fundamentally different from existing approaches because we use peptide groups determined from graph modeling, rather than protein groups or gene groups, as the quantification unit. Using the iPSC and HCC datasets, we demonstrate the ability of our approach in addressing the key limitations of the parsimony-based methods, providing more comprehensive proteome characterization, identifying hundreds of isoform-level regulation events, and enabling streamlined cross-study comparisons.

## Results

### Assessing the challenge and opportunities of isoform characterization

Among the 19,449 protein coding genes annotated in the RefSeq database, 14,698 (75.6%) have more than one protein isoforms, and 3,409 (17.5%) have 10 or more protein isoforms (**Fig. 1a**). Most of isoforms from the same gene have very high sequence similarity (>90%, **Fig. 1b**), highlighting the challenge in discriminating isoforms in shotgun proteomics experiments. However, among the 11,809 genes with three or more protein isoforms, 6,165 (52.2%) have at least one pair of isoforms with a sequence similarity lower than 90%, or an average of one amino acid difference in every 10 amino acids, suggesting the possibility to identify isoform-discriminating peptide sequences for a substantial number of genes.

**Fig. 1:**
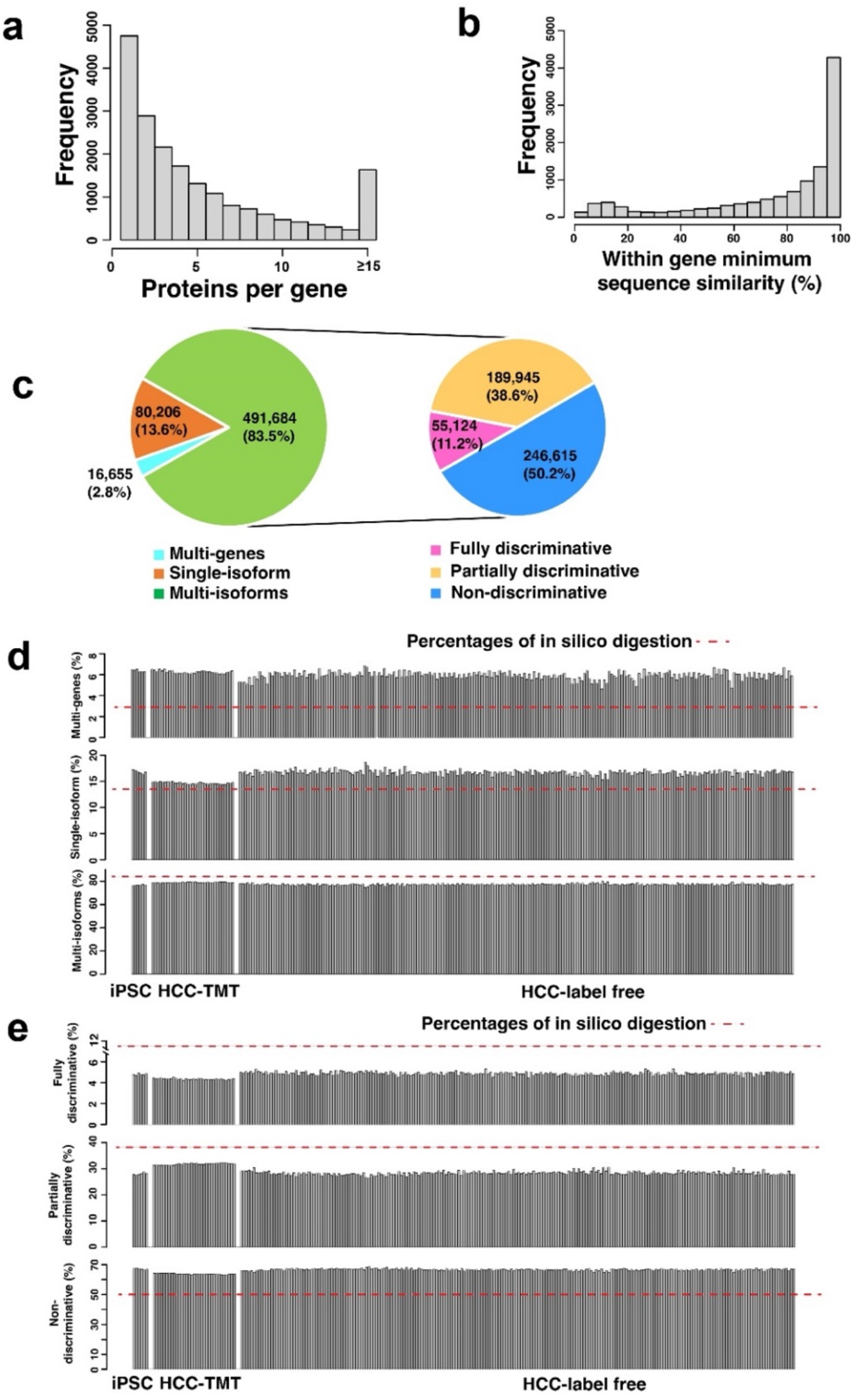
Assessing the challenge and opportunities of protein isoform characterization. **a** Distribution of the number of protein isoforms for all coding genes. **b** Distribution of the minimum within-gene protein isoform sequence similarity. **c** Classification of the *in silico* digested peptides based on their mapping to genes and protein isoforms. **d-e** Classification of the experimentally identified peptides in individual samples in an iPSC cell line study and two hepatocellular carcinoma (HCC) studies.

To further assess the challenge and opportunities of isoform characterization using shotgun proteomics, we performed *in silico* trypsin digestion of the RefSeq protein database to generate fully tryptic peptides with length 7 to 50 and no missed cleavage. Among the 1,883,206 resulting peptide sequences, 2.8% could be associated to multiple genes (*i*.*e*., multi-gene peptides), 13.6% to genes with a single protein isoform (*i*.*e*., single isoform peptides), and 83.5% to genes with more than one isoform (*i*.*e*., multi-isoform peptides). Within the group of multi-isoform peptides, around half could be mapped to all protein isoforms of a gene, and thus providing no information for isoform discrimination (*i*.*e*., non-discriminative peptides); however, another half, or 246,615 peptides, could be uniquely mapped to one isoform (*i*.*e*., fully discriminative peptides) or a subset of isoforms (*i*.*e*., partially discriminative peptides) (**Fig. 1c**).

Next, we compared our *in silico* digestion results with experimental data from a tandem mass tag (TMT)-based iPSC study^21^ and two human HCC studies^22,23^, one TMT based (HCC-TMT) and one label free (HCC-label free). Although two missed cleavage sites were allowed in database searching, less than 5% of identified peptides had missed cleavage sites (**Supplementary Table 1**). Around 6% of the peptides identified in these studies were multi-gene peptides, and the ratios more than doubled the 2.8% estimate from the *in silico* digestion (**Fig. 1d**). This could be explained by possibly higher abundance of these peptides because they can be derived from multiple genes. Percentages of the single isoform peptides in these studies were slightly higher than that in *in silico* digestion, whereas an opposite trend was found for multi-isoform peptides (**Fig. 1d**), suggesting competition among multiple isoforms may reduce transcriptional and/or translational efficiency. Within the group of multi-isoform peptides, the ratios of non-discriminative peptides were about 15% higher in these studies than those in *in silico* digestion, likely due to contributions from multiple isoforms (**Fig. 1e**). Percentages of the fully discriminative peptides and partially discriminative peptides in these studies were lower than those in *in silico* digestion, however, they still accounted for about 35% of the multi-isoform peptides, or about 27% of all identified peptides (**Fig. 1e**).

In summary, experimental data are largely in consistent with the *in silico* digestion results, and both suggest that despite intrinsic challenges, there are a substantial fraction of peptides that hold important information for isoform characterization in shotgun proteomics.

### Tripartite graph modeling of peptides identified by shotgun proteomics

Peptides shared by multiple genes or multiple protein isoforms of the same gene complicate protein inference and quantification. Based on Occam’s razor or the principle of parsimony, the current best practice in the proteomics field is to collapse proteins with the same or subset of supporting peptides into a minimal list of protein groups, and for quantitative rollup, peptides shared by multiple proteins are assigned only to the ones with the most identification evidence. Although practically useful, this parsimonious approach greatly limits the potential for protein isoform characterization. We propose a tripartite graph modeling approach to represent the data more accurately.

The tripartite graph modeling approach involves four major steps (**Fig. 2a-d**). First, a tripartite graph is built with three sets of vertices representing all peptides identified in a study (Pep1-Pep12), proteins to which the peptides can be mapped (Pro1.1-Pro3.1), and host genes of the proteins (Gene1-Gene3), respectively, and the vertices are connected by edges indicating their mapping relationships (**Fig. 2a**). Second, using a graph theory based technique^24,25^, peptides connected to exactly the same set of protein vertices are grouped together and defined as a group of structurally equivalent peptides (SEPEP), leading to eight SEPEPs in **Fig. 2b**. Third, the picked FDR (false discovery rate) approach^26^ is used to estimate target-decoy based FDR at the SEPEP level. Specifically, a SEPEP is considered a target hit if the highest scoring peptide in the SEPEP is from a forward protein sequence, and a decoy hit if the highest scoring peptide is from a decoy protein sequence. SEPEPs with FDR >0.01 are excluded from further analysis (**Fig. 1c**). Finally, the remaining SEPEPs are classified into 5 classes based on their patterns of connections to source proteins and genes in the tripartite graph (**Fig. 2d**). Class 1 through 5 correspond to single isoform SEPEPs, fully discriminative SEPEPs, partially discriminative SEPEPs, non-discriminative SEPEPs, and multi-gene SEPEPs, respectively. Class 1 to 4 SEPEPs are labeled with gene name followed by SEPEP order within the gene and SEPEP class type, *e*.*g*., Gene1_SEPEP.1_C1. Class 5 SEPEPs are labeled with “Multiple” followed by SEPEP order across the whole study and C5, *e*.*g*., Multiple_SEPEP.1_C5. Each SEPEP is also identified by associated gene(s) and protein(s), both alphabetically sorted and concatenated, in a mapping table, which will allow streamlined cross study comparison.

**Fig. 2:**
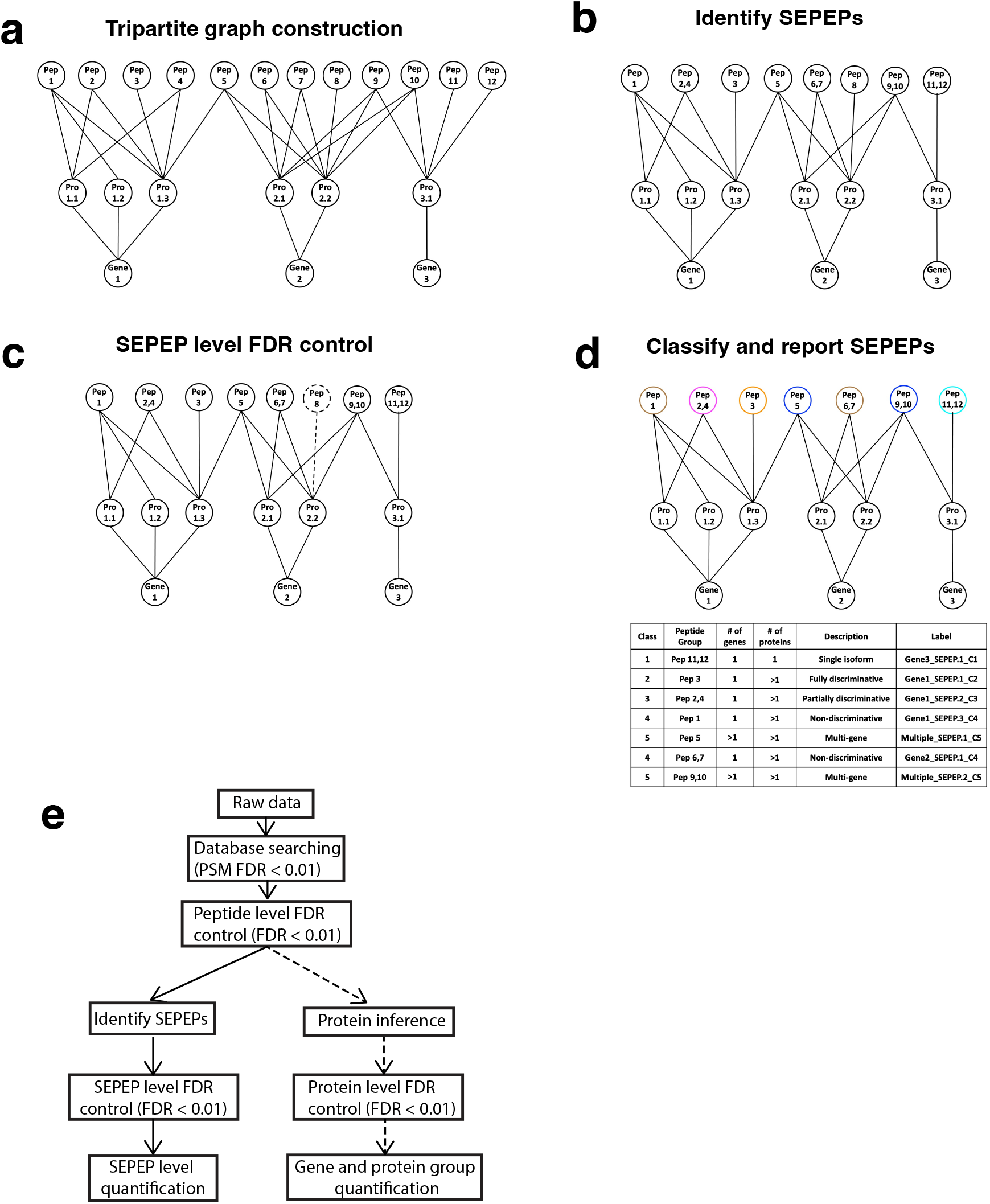
Overview of the tripartite graph modeling approach. **a** Tripartite graph construction: connect peptides to all proteins that contain them, and proteins to the host genes, to form a tripartite graph. **b** SEPEP identification: identify and group structurally equivalent peptides (SEPEP), i.e., peptides connecting to the same set of proteins in the graph. **c** FDR control: calculate SEPEP level FDR and remove SEPEPs with an FDR higher than the pre-specified threshold, e.g., FDR > 0.01. **d** SEPEP classification and reporting: classify SEPEPs based on their patterns of connections and report SEPEP level quantification. **e** A comparison between SEPEP analysis procedure and the classical parsimony protein inference-based procedure.

In contrast to existing methods that make protein inference and then use protein groups or gene groups as the reporting and quantification units, our approach uses SEPEPs as the reporting and quantification units (**Fig. 2e**). All methods share the same database searching, peptide-spectrum match (PSM) FDR control, and peptide FDR control protocols. SEPEP identification and SEPEP level FDR control are performed in parallel to the standard protein inference and protein level FDR control. Finally, the same algorithm can be used to report quantification at SEPEP, gene group, and protein group levels. Specifically, each quantification unit (protein group, gene group, or SEPEP) is quantified based on abundance of all associated peptides using an appropriate method, such as mean, median, or sum, according to the nature of proteomics experiment. SEPEP based quantification is referred to as SEPepQuant.

### SEPepQuant enables more comprehensive proteome characterization

We applied SEPepQuant to the iPSC, HCC-TMT, and HCC-label free data sets mentioned above. The numbers of identified SEPEPs ranged from about 10,000 to 25,000 for each sample or TMT plex, driving by the depth of the proteomics studies (**Fig. 3a**). Although the numbers of identified SEPEPs varied greatly across different data sets, the percentages of SEPEPs failing 1% FDR filtering were around 15% for most samples, except for 11 samples from the HCC-label free data set (**Supplementary Fig. 1a**). About 90% of the rejected SEPEPs in these data sets contained only a single peptide with a mean peptide number of about 1.04 (**Supplementary Fig. 1b**). Therefore, the rejected SEPEPs may represent less robust identifications. After quality control, there were still about 500 - 3500 class 5 SEPEPs (**Fig. 3b**). These multi-gene peptides are typically removed or assigned to a single gene with the strongest identification evidence in parsimonious protein inference, leading to information loss or potential misinterpretation.

**Fig. 3:**
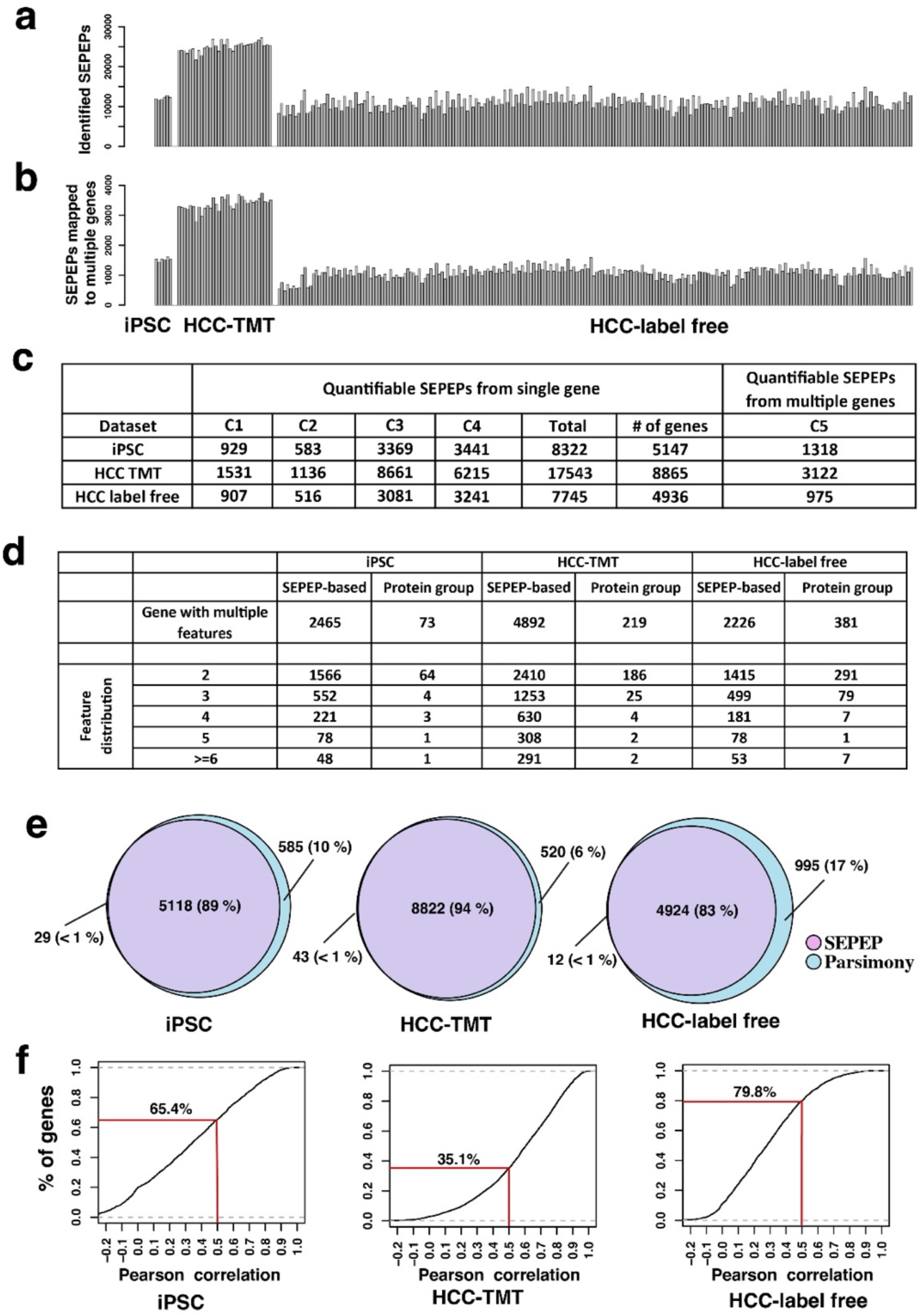
SEPEP-level quality control and quantification in three selected data sets. **a** Numbers of identified SEPEPs. **b** Numbers of multi-gene SEPEPs passing FDR control. **c** Class distribution of the quantifiable SEPEPs. **d** Comparison of numbers of genes with multiple SEPEPs in SEPepQuant analysis and those with multiple protein groups in parsimonious inference. **e** Overlap of genes with quantifiable SEPEPs and quantifiable genes by parsimonious inference. **f** Distribution of the lowest correlations between SEPEPs and host genes with more than one SEPEP.

The SEPepQuant results were further filtered by removing SEPEPs with more than 50% missing values in each data set. SEPEPs passing this criterion were considered as quantifiable SEPEPs. The same criterion was also applied to gene and protein group level quantification derived from parsimonious protein inference. The total number of quantifiable single gene SEPEPs were 8,322, 17,543, and 7,745 from the iPSC, HCC-TMT, and HCC-label free data sets, respectively, and they corresponded to 5,147, 8,865, and 4,936 genes (**Fig. 3c**). Compared with protein group level quantification, which may also provide abundance of multiple distinguishable protein groups for individual genes, SEPepQuant reported 5.8 to 33.8 time more genes with multiple features (**Fig. 3d**). Moreover, SEPepQuant also reported 1,318, 3,122, and 975 multi-genes SEPEPs for the three data sets, respectively, and such information is missing or difficult to track in existing computational tools.

We further compared genes harboring quantifiable SEPEPs with quantifiable genes reported based on parsimonious inference (**Fig. 3e**). The numbers of genes harboring quantifiable SEPEPs were smaller than corresponding gene numbers reported by parsimonious inference across all three data sets. This can be explained by more stringent FDR control caused by the much larger number of candidate SEPEPs compared with candidate genes (**Fig. 3a** and **3d**). Quantifications of the C4 SEPEPs were highly correlated with their host gene quantifications, which is consistent with the observation that more than 60% of peptides from genes with multiple protein isoforms could be mapped to all isoforms (**Supplementary Fig. 1c**). Nevertheless, among genes quantified by both methods and with multiple SEPEPs, many had at least one SEPEP with a correlation less than 0.5 with their host genes, including 65.4%, 35.1%, and 79.8% of the genes from the iPSC, HCC-TMT, and HCC-label free data set, respectively (**Fig. 3f**). Together, these results show that SEPepQuant provides additional resolution that may enable more comprehensive proteome characterization than traditional protein group level or gene level analysis.

### SEPepQuant addresses key limitations of the parsimony-based methods

In parsimonious protein inference, protein isoforms without uniquely identified peptides are largely ignored, and shared peptides are assigned to proteins with the most identification evidence for quantification. Here we use representative examples from the HCC-TMT data set to illustrate the drawback of these simplifications on protein isoform characterization, and the effectiveness of SEPepQuant in addressing these limitations.

*ACTR3* encodes two protein isoforms. The short isoform NP_001264069.1 is translated from a downstream translation initiation site and is shorter at the N-terminus compared to the long isoform NP_005712.1. Among the 17 peptides identified for this gene, three were unique to the long isoform and others were shared between the two isoforms (**Fig. 4a**). Parsimonious protein inference assigned all shared peptides to the long isoform, and the quantification based on all peptides showed that NP_005712.1 was significantly lower in tumors compared with normal adjacent tissues (NATs) (p=1.9e-4). Meanwhile, no quantitative information was provided for the short isoform. In contrast, SEPepQuant reported two SEPEPs. ACTR3_SEPEP.2_C2 was associated with the three NP_005712.1-specific peptides, and it was significantly higher in tumors compared with NATs (p=2.3e-11). ACTR3_SEPEP.1_C4 was associated with the shared peptides, and it was lower in tumors compared with NATs with a marginal significance (p=0.02). Despite the lack of NP_001264069.1-specific peptides, it was not difficult to infer that this isoform was suppressed in tumors based on the strong elevation of NP_005712.1. Thus, SEPepQuant provided useful information for both isoforms, whereas the parsimonious inference provided only incorrect information for the long isoform.

**Fig. 4:**
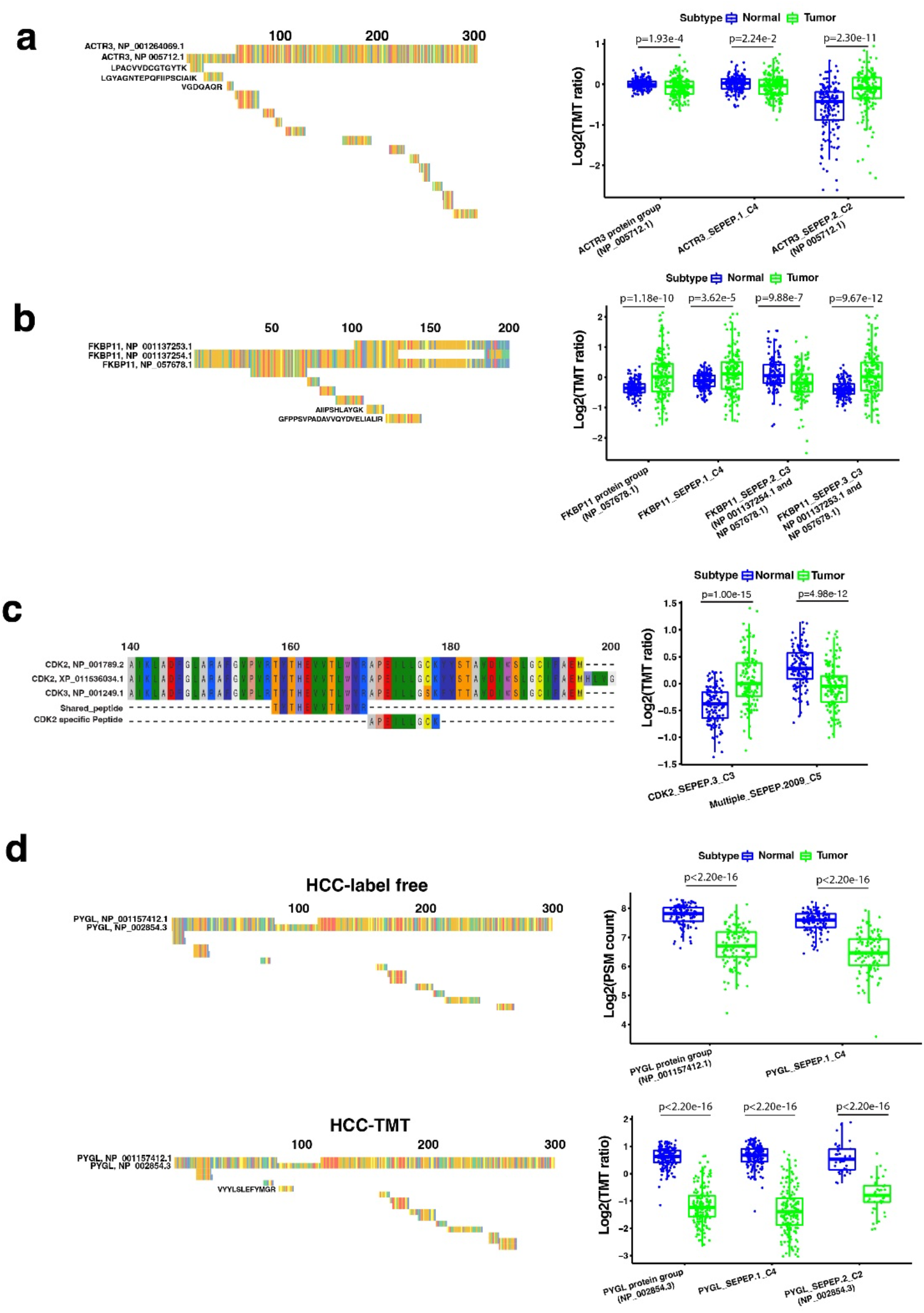
SEPepQuant addresses key limitations of the parsimony-based methods. **a** Identified peptides of ACTR3 from 1-300 bp. Tumor and normal comparisons of ACTR3 SEPEP and protein group levels. **b** Identified peptides of FKBP11 from 0-200 bp. Tumor and normal comparisons of FKBP11 SEPEP and protein group levels. **c** Multi-gene SEPEP provides information on CDK3, a gene without uniquely mapped peptides. **d** protein inference of PYGL in HCC label-free and HCC-TMT data sets. P values were computed based on two sided t-test.

*FKBP11* encodes three protein isoforms, and none of the six peptides identified for this gene could be uniquely mapped to a single isoform (**Fig. 4b**). Because NP_057678.1 could explain all six observed peptides but the other two isoforms could not, parsimonious protein inference assigned all six peptides to NP_057678.1, and the resulting quantification showed that this isoform was significantly increased in tumor samples compared with NATs. SEPepQuant reported quantification results for three SEPEPs. Both the non-discriminative FKBP11_SEPEP.1_C4 (one peptide) and the partially discriminative FKBP11_SEPEP.3_C3 (NP_001137253.1 and NP_057678.1, one peptide) had higher abundance in tumors, whereas the partially discriminative FKBP11_SEPEP.2_C3 (NP_001137254.1 and NP_057678.1, four peptides) had lower abundance in tumors. Although the change of NP_057678.1 remained difficult to determine, SEPepQuant results clearly suggested decreased abundance of NP 001137254.1 and increased abundance of NP_001137253.1 in tumor samples.

In addition to providing higher resolution for characterizing multiple protein isoforms encoded by the same gene, SEPepQuant also reports quantifications for multi-gene SEPEPs (C5 SEPEPs), which may provide useful information that is typically missed in the parsimony-based methods. For example, Multiple_SEPEP.2009_C5 was associated with a peptide that could be mapped to two CDK2 isoforms and one CDK3 isoform (**Fig. 4c**). In parsimonious inference, because the two CDK2 isoforms were also supported by another CDK2-specific peptide, this shared peptide was assigned to the CDK2 isoforms and thus CDK3 was not quantified. With SEPepQuant, Multiple_SEPEP.2009_C5 was reported to be highly significantly decreased in tumors compared with NATs (p=4.96e-12). Although this SEPEP was associated with two CDK2 isoforms and one CDK3 isoform, it was possible to associate the decrease of this SEPEP specifically to the CDK3 isoform because another SEPEP (CDK2_SEPEP.3_C3) uniquely mapped to the two CDK2 isoforms were highly significantly overexpressed in tumors (p=1.00e-15).

Parsimonious inference may also complicate cross-study comparisons. For example, *PYGL* encodes two protein isoforms differing in an exon skipping event (**Fig. 4d**). The HCC-TMT study identified many shared peptides and one unique to the long isoform. Accordingly, all peptides were assigned to the long isoform for reporting. In the closely related HCC-label free study, all identified peptides were shared between the two isoforms. According to the parsimonious principle, all peptides were assigned to the short isoform for reporting. Although significantly decreased expression of PYGL in tumors compared with NATs was observed in both studies, and most of the identified peptides were identical in the two studies (**Fig. 4d**), different protein isoforms reported by the two studies could cause confusions in a cross-study comparison. In addition to reporting one SEPEP associated with the long isoform-specific peptide in the HCC-TMT study, SEPepQuant also reported a non-discriminative SEPEP, which was consistently decreased in tumors compared with NATs in both studies, which helped eliminate potential confusion.

### Protein isoform expression changes during iPSC differentiation into cardiomyocyte

To demonstrate the practical utility of SEPepQuant, we compared SEPepQuant analysis results of the iPSC dataset with those from gene level quantifications reported by FragPipe. The iPSC dataset was generated by TMT-based shotgun proteomics on iPSC cells cultured over 14 days and harvested daily^21^. As a positive control, we checked the gene and SEPEP level results of TPM1, which was found to have two regulated protein isoforms in the original study. TPM1 showed significant up-regulation with culture time based on gene-level quantification by FragPipe (**Supplementary Fig. 2a**). Consistent with the original study, SEPepQuant identified a group of isoforms recognized by TPM1_SEPEP.2_C3, which differed from the canonical TPM1 sequence at residues 189-212 by mutually exclusive exon (MXE) splicing and were up-regulated from day 0 to day 7 and then down-regulated from day 7 to day 14 (**Supplementary Fig. 2b**), as wells as another group of isoforms recognized by TPM1_SEPEP.8_C3, which differed from the canonical TPM1 sequence at residues 41-80 by MXE splicing and were upregulated at day14 compared with day7 (**Supplementary Fig. 2c**). In addition, SEPepQuant further identified another down-regulated isoform group recognized by TPM1_SEPEP.6_C3, which used an alternative translation start site (**Supplementary Fig. 2d**). Thus, SEPepQuant not only confirmed existing findings but also revealed new information.

Next, we correlated quantifiable genes and SEPEPs with culture time to identify genes and SEPEPs showing monotonic abundance changes during iPSC differentiation into cardiomyocyte. Among the 2028 quantifiable genes with multiple SEPEPs, most showed concordant alterations at gene and SEPEP levels (**Fig. 5a** and **Supplementary Table 2**). However, 152 SEPEPs from 136 genes showed a significant positive correlation with culture time without matching significance at the gene level, including 10 SEPEPs from nine genes showing a significant negative correlation at the gene level. As an example, PES1_SEPEP.2_C3 was significantly up regulated with culture time, but PES1_SEPEP.1_C4 and the gene level quantification were significantly down regulated (**Fig. 5b**). Similarly, 122 SEPEPs from 110 genes showed significant negative correlation with culture time without matching significance at the gene level, including three SEPEPs from three genes showing a significant positive correlation at the gene level. For example, DPYSL3_SEPEP.2_C2, which included a single protein NP_001184223.1 and with 5 unique identified peptides (**Supplementary Fig. 2e**), was significantly down regulated with time; but both DPYSL3_SEPEP.1_C4 and the gene level quantification showed a pattern of down regulation from day 0 to day 7 and upregulation from day 7 to day 14 (**Fig. 5c**).

**Fig. 5:**
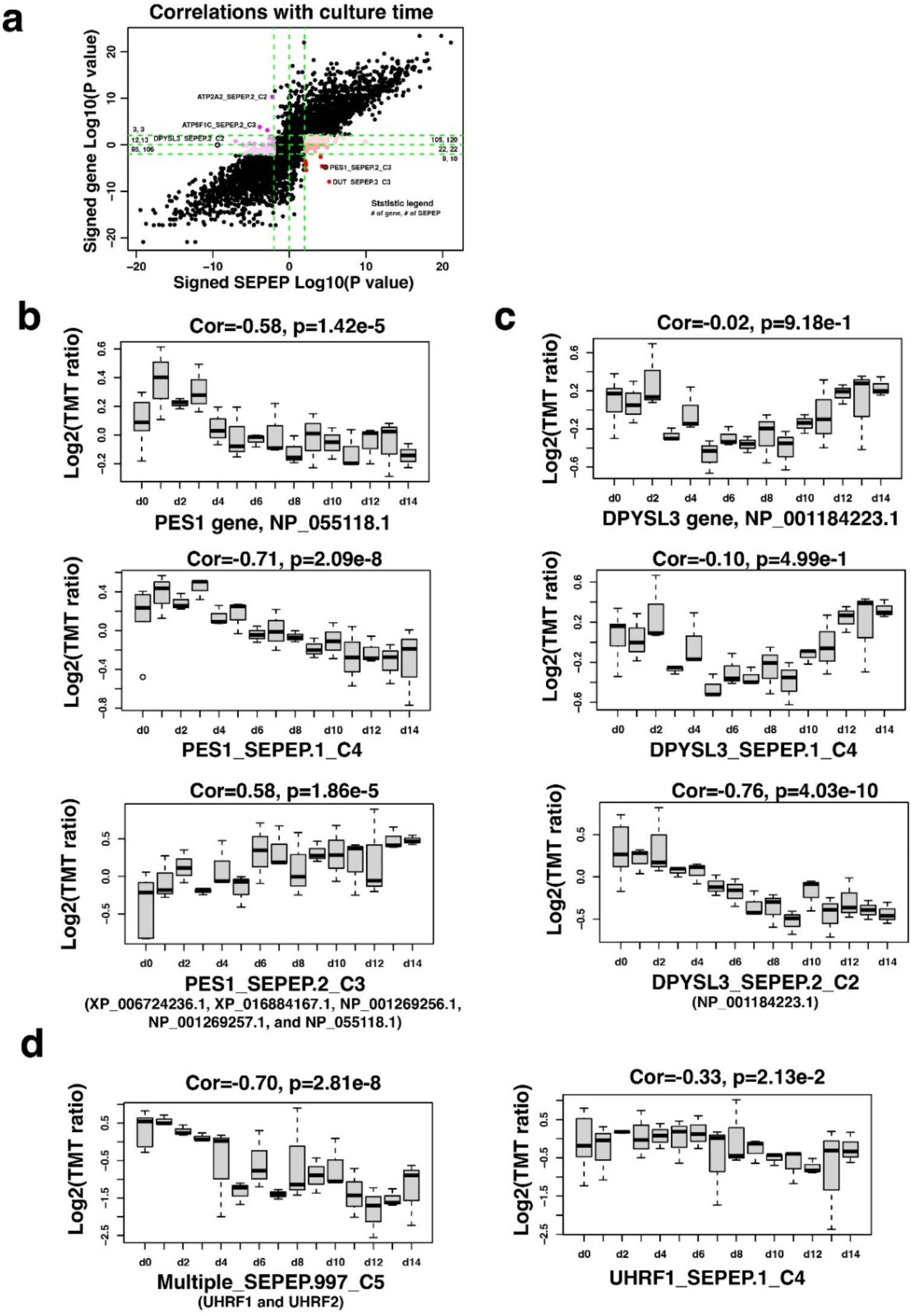
Application of SEPepQuant to an iPSC data set. **a** Scatter plot comparing time-dependent regulations at gene and SEPEP levels. **b** Time-dependent regulation of PES1 at gene and SEPEP levels. **c** Time-dependent regulation of DPYSL3 at gene and SEPEP levels. **d** Time-dependent regulation of a multi-gene SEPEP Multiple_ SEPEP.997_C5, which is associated with both UHRF1 and UHRF2, and a single-gene SEPEP UHRF1_ SEPEP.1_C4, which is specific to UHRF1.

Our analysis also identified 221 multi-gene SEPEPs showing significant correlation with culture time (**Supplementary Fig. 2f**). The most significantly positively correlated multi-gene SEPEP, Multiple_SEPEP.122_C5, was associated with four calmodulin-dependent protein kinase genes *CAMK2D, CAMK2A, CAMK2B*, and *CAMK2G*. The most significantly negatively correlated multi-gene SEPEP Multiple_SEPEP.978_C5 was associated with four zinc finger genes *ZNF93, ZNF431, ZNF714*, and *ZNF92*. Although it is difficult to attribute the associations to specific genes, these observations still revealed important roles of these gene families in iPSC differentiation. Moreover, data specific to some genes in a multi-gene SEPEP could be leveraged to improve the interpretation of the association observed for the multi-gene SEPEP. For example, the SEPEP Multiple_SEPEP.997_C5 was associated with both *UHRF1* and *UHRF2*, and it was found to be significantly negatively correlated with time (**Fig. 5d**). Because UHRF1_SEPEP.1_C1 was not significantly correlated with time (**Fig. 5d**), it is logical to infer that the significant negative association between Multiple_SEPEP.997_C5 and time was driven primarily by *UHRF2* even though it was not specifically quantified in this data set.

### Protein isoforms associated with liver cancer development and prognosis

The HCC-TMT dataset included liver tumor samples and paired NATs from 165 patients, and overall survival information is also available for these patients. We performed tumor vs NAT comparison and survival analysis based on SEPepQuant and gene-level quantifications reported by FragPipe, respectively. In the tumor vs NAT comparison, most of the 4,481 genes with multiple SEPEPs showed concordant alterations at gene and SEPEP levels (**Fig. 6a, Supplementary Table 3**). However, 405 SEPEPs from 336 genes showed a significant increase in tumor samples without matching significance at the gene level, including 82 SEPEPs from 73 genes showing a significant decrease in tumor samples at the gene level. Similarly, 406 SEPEPs from 342 genes showed a significant decrease in tumor samples without matching significance at the gene level, including 98 SEPEPs from 89 genes showing a significant increase in tumor samples at the gene level. In the survival analysis, a higher level of concordance was observed between SEPEP level and gene level results (**Fig. 6b, Supplementary Table 4**). However, there were still hundreds of SEPEPs showing significant positive or negative associations with survival without matching significance at the gene level. Notably, 19 genes showed significantly increased expression in tumors compared with normal samples as well as significant association with poor prognosis at the SEPEP level, and these associations were not observable at the gene level (**Fig. 6c, Supplementary Table 5**).

**Fig. 6:**
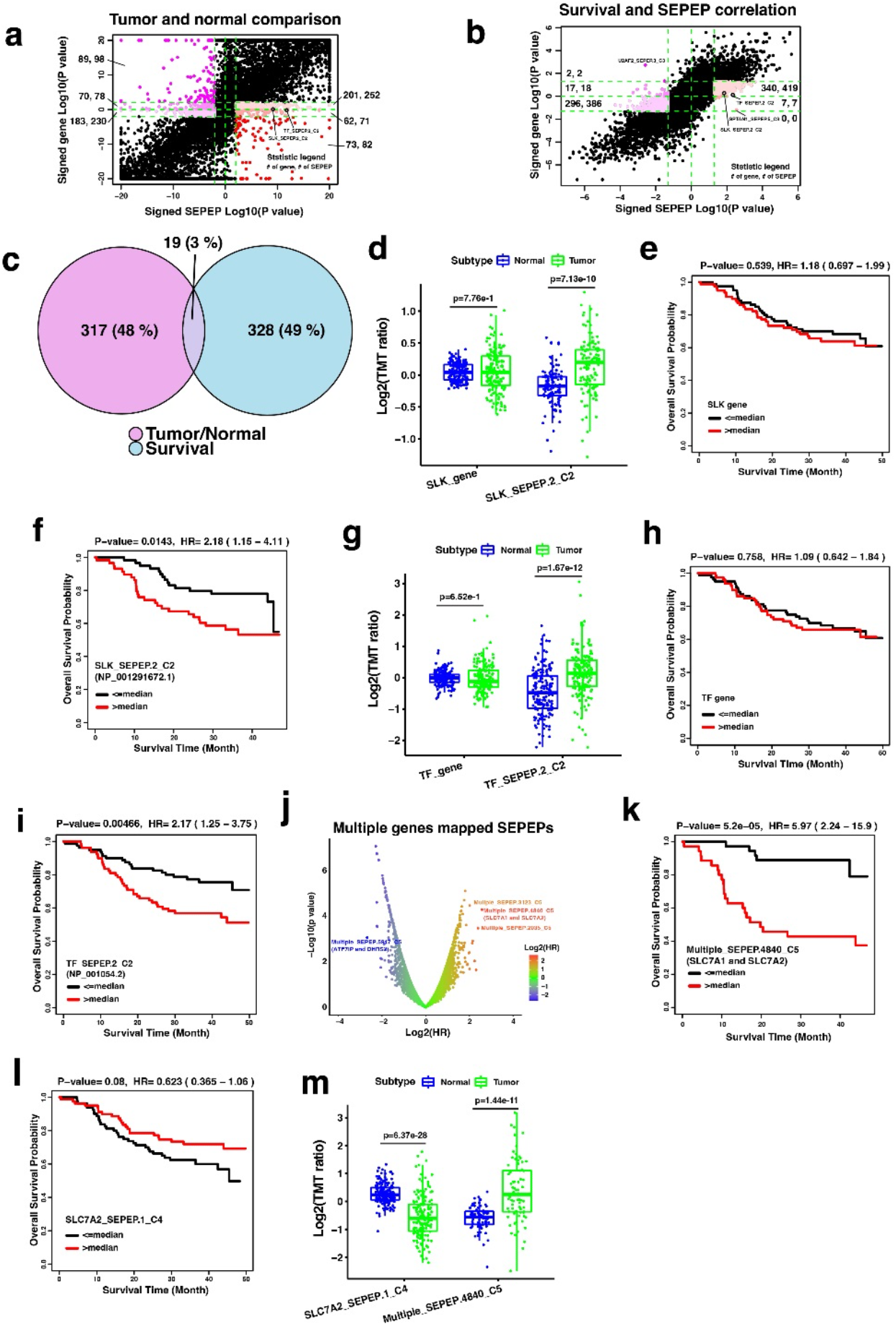
Application of SEPepQuant to an HCC-TMT data set. **a** Scatter plot of tumor versus normal comparison results at gene and SEPEP levels. **b** Scatter plot of survival association results at gene and SEPEP levels. **c** Overlap of genes showing significantly increased expression in tumors compared with normal samples as well as significant association with poor prognosis at the SEPEP level but not at the gene level. **d** Tumor versus normal comparisons based on SLK gene and SLK_SEPEP.2_C2 abundance, respectively. **e-f** Kaplan-Meier plots comparing overall survival for patients stratified by the median SLK gene level abundance and median SLK_SEPEP.2_C2 abundance, respectively. **g** Tumor versus normal comparisons based on TF gene and TF_SEPEP.2_C2 abundance, respectively. **h-i** Kaplan-Meier plots comparing overall survival for patients stratified by the median TF gene level abundance and median TF_SEPEP.2_C2 abundance, respectively. **j** Associations between survival and multi-genes SEPEPs. **k-l** Kaplan-Meier plots comparing overall survival for patients stratified by the median Multiple_SEPEP.4840_C5 abundance and median SLC7A2_SEPEP.1_C4 abundance, respectively. **m** Tumor versus normal comparisons based on Multiple_SEPEP.4840_C5 and SLC7A2_SEPEP.1_C4 abundance, respectively.

One of the 19 was SLK_SEPEP.2_C2, a fully discriminative SEPEP associated with a single protein isoform NP_001291672.1 encoded by the STE20-like serine/threonine-protein kinase gene *SLK* (**Fig. 6d-f**). Identification and quantification of this SEPEP were based on two junction peptides specific to NP_001291672.1 (**Supplementary Fig. 3a**). Among the three protein isoforms encoded by *SLK*, SEPepQuant connected this specific isoform to liver cancer development and prognosis, suggesting a critical pro-tumor role of the exon skipping event. Although the quantification for the NP_001291672.1-specific SEPEP was very sparse in the independent HCC-label free data set, samples with identification of NP_001291672.1 specific peptide KKEEQEFVQK had significantly worse survival compared with samples without identification of this peptide (**Supplementary Fig. 3b**), which provided independent confirmation of our finding in the TMT data set. Protein group-level quantification from FragPipe reported two protein groups with representative proteins NP_001291672.1 and NP_055535.2, respectively. However, the protein group represented by NP_001291672.1 showed no significant association with survival in FragPipe quantification (**Supplementary Fig. 3c**). This is because multiple shared peptides were assigned to this protein group based on the parsimony principal, but shared peptides derived from the other protein group, which had significant association with good prognosis (**Supplementary Fig. 3d)**, could greatly dilute the signal specific to NP_001291672.1.

Another SEPEP showing significantly increased expression in tumors compared with normal samples as well as significant association with poor prognosis but without concordant changes at the gene level was TF_SEPEP.2_C2, a fully discriminative SEPEP associated with a single protein isoform NP_001054.2 encoded by the transferrin (*TF*) gene (**Fig. 6g-i**). Identification and quantification of this SEPEP were based on three peptides mapping to the N-terminal region specific to NP_001054.2 (**Supplementary Fig. 3e**). Among the three protein isoforms encoded by the *TF* gene, SEPepQuant connected this specific isoform to liver cancer development and prognosis. Despite sparse identification of the isoform-specific peptides, the trend was confirmed in the independent HCC label-free data set (**Supplementary Fig. 3f-g**). Protein group-level report from FragPipe only reported one protein group with NP_001054.2 as the representative protein because it can explain all identified peptides (**Supplementary Fig. 3e**). However, quantification of this protein group based on all peptides was equivalent to gene-level quantification, which showed no significant difference in tumor vs normal comparison and no significant association in survival analysis (**Fig. 6g-h**).

Our analysis also identified 423 and 418 multi-gene SEPEPs with a significant association with good or poor prognosis, respectively (**Fig. 6j**). Among them, multiple_SEPEP.5947_C5 showed the lowest hazard ratio. This SEPEP was associated with two genes *ATF7IP* and *DHRS2*, and DHRS2 has been reported to inhibit cell growth and motility in esophageal squamous cell carcinoma ^27^. Multiple_SEPEP.4840_C5 showed the strongest association with poor prognosis. This SEPEP was associated with two genes *SLC7A1* and *SLC7A2* (**Fig. 6k**). Interestingly, SLC7A2_SEPEP.1_C4 was associated with slightly longer patient survival (**Fig. 6l**), suggesting that the association between Multiple_SEPEP.4840_C5 and poor prognosis was driven by SLC7A1. Consistent with this inference, Multiple_SEPEP.4840_C5 showed significantly higher abundance in tumors compared with normal samples, but SLC7A2_SEPEP.1_C4 was significantly decreased in tumors. Despite the lack of detection of gene-specific peptides for *SLC7A1*, SEPepQuant results clearly suggested a pro-tumor role of this gene, which was previously reported in ovarian cancer^28^.

## Discussion

Shotgun proteomics has become an essential tool for protein identification and quantification in biomedical research. However, shotgun proteomics-based protein isoform identification and quantification remains an open challenge, hampering a thorough understanding of protein isoform regulation and their roles in normal and disease biology. We developed SEPepQuant, a graph theory-based approach that uses groups of structurally equivalent peptides in a peptide-protein-gene tripartite graph, instead of protein groups or gene groups, as the identification and quantification unit to enable comprehensive protein isoform characterization in shotgun proteomics. In three experimental data sets, SEPepQuant identified 5.8-33.8 time more genes with multiple quantification units compared with that by parsimony-based protein inference (**Fig. 3**). For genes with multiple SEPEPs, 35.1% - 79.8% had at least one SEPEP with a below 0.5 correlation with the corresponding gene abundance, suggesting extensive isoform-specific regulation. Indeed, analysis based on SEPepQuant quantification results revealed more than 100 genes with protein isoform-level regulation during cardiomyocyte differentiation and hundreds of protein isoform level regulatory events with significant associations to liver cancer development and prognosis.

Parsimony-based protein inference was introduced at the early stage of proteomics research to address the problem of overreporting the number of identified proteins in shotgun proteomics studies^16,17^, and it has since become the dominant method in the field. However, the consequence of this method on protein quantification has not been formally evaluated. Our analysis in this paper revealed several key limitations associated with the parsimony-based methods, including ignoring protein isoforms and genes without uniquely identified peptides, incorrect or inaccurate quantification of protein isoforms by simply assigning shared peptides to isoforms with the largest number of identified peptides, and complicating cross-study comparisons because of different reporting isoform selection driven by minor changes in peptide detection in different studies. We showed that SEPepQuant is able to address these limitations, leading to more comprehensive and accurate analysis of protein isoforms.

SEPepQuant identified hundreds of protein isoform level regulatory events from both the iPSC and liver cancer data sets, highlighting widespread impact of protein isoform level regulation in normal and disease processes. Notably, 19 genes showed significantly increased expression in liver tumors compared with normal samples as well as significant association with poor prognosis at the SEPEP level but not at the gene level. Among these, *SLK* encodes a kinase that promotes apoptosis^29^. The pro-tumor SEPEP of *SLK* is a fully discriminative SEPEP and thus the pro-tumor effect could be attributed to the associated protein isoform NP_001291672.1. Compared to the longer *SLK* isoform NP_055535.2, this short isoform has a skipped exon, which encodes a section of a coiled-coil domain that mediates homodimerization to enhance SLK activity^30^. Therefore, this exon skipping may lead to reduced SLK activity and decreased apoptosis to facilitate tumor progression. Consistently, a recent analysis of RNASeq data from melanoma tumors has shown that expression of the long isoform is decreased whereas the short isoform is increased in metastatic tumors compared with primary tumors, suggesting a role of the exon skipping in facilitating metastasis^31^. Transferrin is another gene found to be correlated with poor prognosis at the SEPEP level but not gene level. This pro-tumor SEPEP is also a fully discriminative SEPEP associated with NP_001054.2, the longest isoform of the transferrin gene. Transferrin is synthesized primarily in liver and secreted into serum, with a half-life of eight days in the serum^32^. The unique N-terminal sequence of this isoform and the long half-life of the protein make it a promising candidate of serum biomarker of liver cancer prognosis for further investigation.

In summary, our analysis provides strong evidence to support a critical and widespread role of protein isoform regulation in normal and disease processes, and SEPepQuant is expected to have broad applications to biological and translational research to boost scientific discoveries.

## Methods

### Protein database

RefSeq gene annotation and matched protein database with all NP, XP, and YP protein sequences were selected for this study (downloaded on 06/28/2020)^33^.

### In-silico digestion

Protein sequences were cut after any K and R except followed by P using in house script. This corresponds to a full tryptic digestion with no missed cleavage site^34,35^. Resulted peptide sequences with length <7bp or >50bp were excluded from our analysis.

### Protein sequence similarity

Protein sequences for genes with multiple protein isoforms were aligned using clustalw2 with default parameters^36^. Then, clustalo was applied to above multiple sequence alignments to calculate sequence similarity with percent identities as sequence distance^37^. Pair-wise sequence similarities were extracted from the clustalo percentage identity matrix.

### Experimental data sets

Three published data sets were selected to cover different sample types and proteomics platforms. The iPSC data set is a human cell line data set generated on a 10-plex TMT platform. In this study, the human iPSC cells were cultured over 14 days and harvested daily for proteomics analyzing^21^. The HCC TMT data set includes 165 tumor samples from hepatocellular carcinoma patients and matched adjacent normal tissues, and the data were generated on a 11-plex TMT plagrom^22^. The HCC label-free data set includes 111 tumor samples from hepatocellular carcinoma patients and matched adjacent normal tissues, and the data were generated on a label-free platform^23^.

### Database searching and protein quantification using FragPipe

The comprehensive shotgun proteomics analysis pipeline FragPipe (https://fragpipe.nesvilab.org/; V17.0) with MSFragger (v3.4) and philosopher (v4.2.1) was used for database searching and protein group level and gene level quantification for all three data sets^14,15^. The same parameters and modifications from the original publication were used for database searching. PeptideProphet and ProteinProphet with default parameters were used for peptide validation and protein inference^38,39^. PSM count was used for label free data quantification, and TMT ratio was used for quantification in TMT-based data sets. Median centering was used as the normalization methods for TMT-based data sets.

### SEPEP quantification and normalization

For label free data, the sum of PSMs of all peptides from a SEPEP was used to quantify the SEPEP. For TMT-based data, the median value of the TMT ratios of all peptides from a SEPEP was used to quantify the SEPEP. Normalization factors derived from median centering of the gene level analysis described above was used to normalize SEPEP quantifications.

## Supporting information

Manuscript-supplemental

## Data Availability

The human iPSC^21^ and HCC-label free^23^ data sets were downloaded from PRIDE (www.ebi.ac.uk/pride) with accession numbers PXD013426 and PXD006512. The HCC-TMT^22^ proteome data was downloaded from NODE (https://www.biosino.org/node) by accession number OEP000321. Processed quantification data and downstream analysis results were included Supplementary Tables 1-14.

## Code Availability

Code for SEPepQuant analysis can be accessed at https://github.com/bzhanglab/SEPepQuant.

## Contributions

Y.D. and B.Z. conceived the project and designed the study. Y.D. processed proteomics data and performed the analyses with help from B.Z, X.Y., and L.K.O.. Y.D. and B.Z. wrote the manuscript. All authors edited and approved the manuscript.

## Acknowledgements

This study was supported by grants U24 CA210954, U24 CA271076, and R01 CA245903 from the National Cancer Institute (NCI), the Cancer Prevention & Research Institutes of Texas (CPRIT) award RR160027, and funding from the McNair Medical Institute at The Robert and Janice McNair Foundation. B.Z. is a CPRIT Scholar in Cancer Research and a McNair Scholar.

## Supplementary Information

Supplemental figure 1: SEPEP level quality control

Supplemental figure 2: Evaluation of SEPepQuant on an iPSC data set

Supplemental figure 3: Evaluation of SEPepQuant on two liver cancer data sets

Supplementary table 1: Percentage of peptides with missed cleavage site(s)

Supplementary table 2: iPSC-TMT gene and SEPEP correlation with culture time

Supplementary table 3: HCC-TMT gene and SEPEP tumor versus normal comparison

Supplementary table 4: HCC-TMT gene and SEPEP survival analysis

Supplementary table 5: Overlapping significant genes from the HCC-TMT tumor versus normal and survival analyses.

Supplementary table 6: iPSC-TMT SEPEP quantification, median centered

Supplementary table 7: iPSC-TMT SEPEP mapping table

Supplementary table 8: iPSC-FragPipe quantification

Supplementary table 9: HCC-TMT SEPEP quantification, median centered

Supplementary table 10: HCC-TMT SEPEP mapping table

Supplementary table 11: HCC-TMT FragPipe quantification

Supplementary table 12: HCC-label free SEPEP PSM count

Supplementary table 13: HCC-label free SEPEP mapping table

Supplementary table 14: HCC-label free FragPipe quantification

