## Supplementary material for "SEPepQuant enables comprehensive protein isoform characterization in shotgun proteomics": Manuscript-supplemental

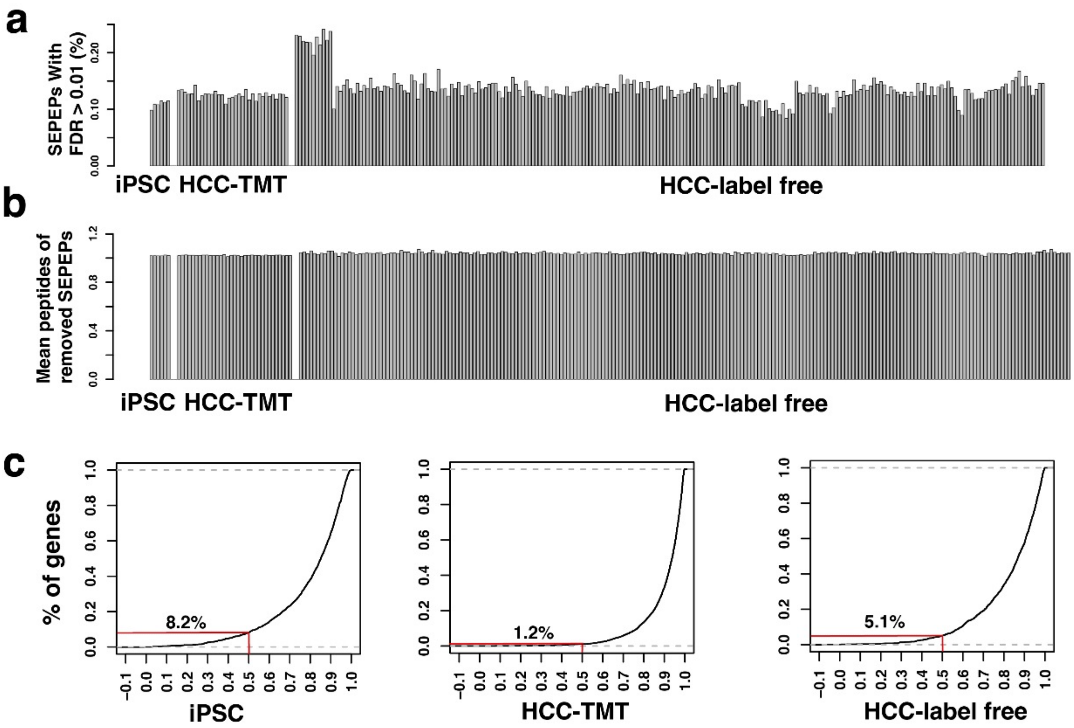


**Supplemental figure 1: SEPEP level quality control**. (a) Percentages of SEPEPs with FDR >0.01. (b) Mean peptide numbers of SEPEPs with FDR >0.01. (c) Distributions of correlations between FragPipe gene level and SEPEP C4 quantification.

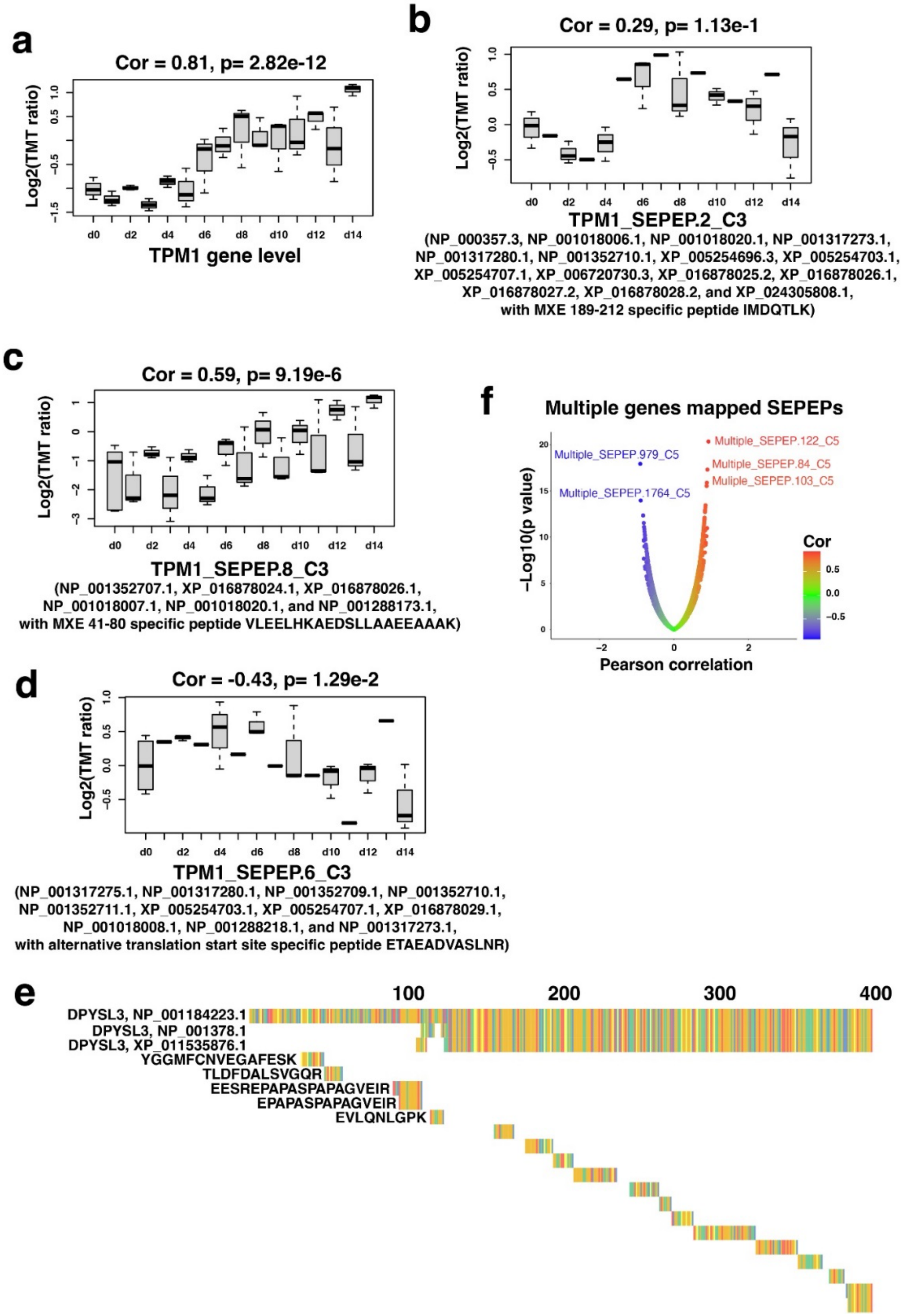


**Supplemental figure 2: Evaluation of SEPepQuant on an iPSC data set.** (a-d) Protein abundance and culture time correlations of TPM1 gene and selected SEPEPs. (e) Identified peptides of DPYSL3 on iPSC data set from 1- 400bp. (f) Correlations between cell culture time and multi-genes SEPEPs.

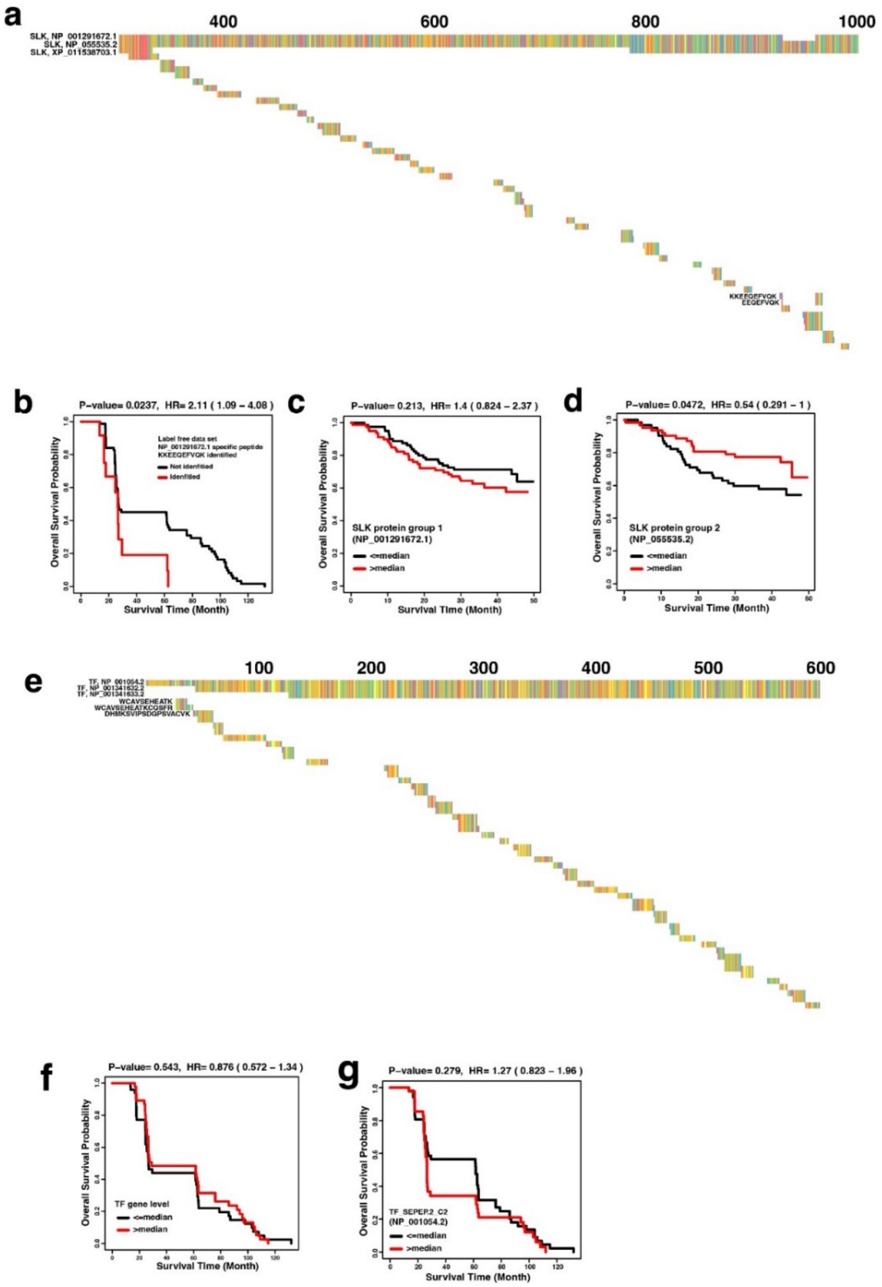


**Supplemental figure 3: Evaluation of SEPepQuant on two liver cancer data sets**. (a) Identified peptides of SLK on HCC-TMT data set. (b) Survival comparison between sample with and without NP_001291672.1 specific peptide KKEEQEFVQK on HCC-label free data set. (c) Correlation of NP_001291672.1 abundance by FragPipe and survival on HCC-TMT data set. (d) Correlation of NP_055535.2 abundance by FragPipe and survival on HCC-TMT data set. (e) Identified peptides of TF on HCC-TMT data set. (f-g) TF gene level and TF_SEPEP.2_C2 abundance and survival correlations on HCC-label free data set.
